# Nano-enhanced optical gene delivery to retinal degenerated mice

**DOI:** 10.1101/509349

**Authors:** Subrata Batabyal, Sivakumar Gajjeraman, Sulagna Bhattacharya, Weldon Wright, Samarendra Mohanty

**Affiliations:** Nanoscope Technologies LLC, 1312 Brown Trail Bedford, Texas, USA, 76022

**Keywords:** Ocular gene therapy, optical delivery, optogenetics, dry-AMD

## Abstract

The efficient and targeted delivery of genes and other impermeable therapeutic molecules into retinal cells is of immense importance for therapy of various visual disorders. Traditional methods for gene delivery require viral transfection, or chemical methods that suffer from one or many drawbacks such as invasiveness, low efficiency, lack of spatially targeted delivery, and can generally have deleterious effects such as unexpected inflammatory responses and immunological reactions. Here, we introduce a continuous wave near-infrared laser-based Nano-enhanced Optical Delivery (NOD) method for spatially controlled delivery of opsin-encoding genes into retina *in-vivo*. In this method, the optical field enhancement by gold nanorods is utilized to transiently permeabilize cell membrane enabling delivery of exogenous impermeable molecules to nanorod-binding cells in laser-irradiated regions. The successful delivery and expression of opsin in targeted retina after in-vivo NOD in the mice models of retinal degeneration opens new vista for re-photosensitizing retina with geographic atrophies as in dry age-related macular degeneration (AMD).

## 1 Introduction

Dry age-related macular degeneration (Dry-AMD) involving dysfunction of retina is characterized by degeneration of photoreceptors. The degree of visual loss increases with ageing and this is a major concern for our demographic changes towards elderly population. The eye offers an excellent target for gene therapy (replacement of defective genes with new functional genes) because it is easily accessible and relatively immune privileged. Most of the current clinical ophthalmic treatments using gene^1^, RNA aptamer^2^ and shRNAs^3^ for therapeutic benefits are utilizing viral vectors, which suffers from drawbacks such as low efficiency, unexpected inflammatory responses, especially innate and immune barriers^4^, toxicity^5^, and potential recombination of or complementation *in vivo* which can generate virulent viral pathogens after vector delivery^6^. In addition, limitation on size of plasmid that can be packaged by the most-commonly used safe (AAV) virus vectors^7,8^ may create hindrance to deliver novel gene-editing large molecules such as CRISPR/Cas9 for visual disorders. Furthermore, the ability to deliver therapeutic genes into spatially targeted higher order retinal neurons of degenerated retina (e.g. geographic atrophic areas of macula in AMD ^9-13^) is advantageous, for example in optogenetic activation therapy. Therefore, there have been significant efforts in developing alternative, non-viral methods such as physical (electroporation^14,15^, ultrasound^16^ etc.) and chemically-mediated (e.g. biodegradable polymer ^17,18^) methods^19,20^. These methods also suffer from one or more drawbacks, such as invasiveness, reduced efficiency, lack of spatially-targeted delivery and/or having several deleterious effects on transfected cells^21^. The advent of optoporation by focused ultrafast near infrared laser microbeam has allowed mitigating the drawbacks of these methods. Recently, we demonstrated use of near IR laser microbeam for delivery of impermeable molecules to neurons ^22^, opsin-encoding genes to cells ^23^ and localized areas of the retina^24^. However, the extreme difficulties in maintaining tight focus inside tissue and requirement of high peak power to create holes in the cell membrane, has limited the use of laser-based delivery to small animal applications^25^. Therefore, there is an imminent need for development and optimization of new and efficient non-viral method that can deliver therapeutic molecules irrespective of their size to spatially targeted regions of retina in a minimally invasive manner.

Recently, optogenetic therapies have been evaluated for vision restoration in mice models of retinal degeneration either by non-specific stimulation of retina^26^ or in a cell-specific manner for retinal ganglion cells^27-31^ and ON bipolar cells^32-33,34^. Further, use of chloride-channel opsin (NpHR) expressing in longer-persisting cone photoreceptor protein has shown promise for restoration of vision^35^. The primary method for delivery of opsin-encoding genes to adult retina employs viral transfection which has high transduction efficiency and persistent gene expression^36^. Both non-specific transfection of the retina as well as promoter-specific targeting of retinal ganglion cells of adult mice and rats using viral methods have been successful^27-30, 32-33, 37-38^. However, clinical translation of optogenetic activation for vision restoration has a major challenge: (i) requirement of external activation light or opsin with significantly higher sensitivity to ambient light; and (ii) method to optogenetically treat only those areas of the retina that have undergone degeneration rather than the whole retina. Effective therapy of visual disorders involving geographic atrophies of the retina requires localized delivery of the targeted molecules to specific retinal cells in atrophied regions.

Herein, we introduce use of nano-enhancement of near-infrared optical field of continuous wave (CW) near-infrared (NIR) laser beam by surface plasmon resonance (SPR) near targeted retinal cells for gene delivery. To allow ambient-light based activation^34^, we utilized multi-characteristic-opsin (MCO) encoding plasmids to sensitize retinal cells in degenerated retina. It would be highly desirable to re-induce transgene expression by reinjection, and NOD provides this opportunity.

## 2 Materials and Methods

### Ethics statement

All experimental procedures were conducted according to the Nanoscope Technologies-Institutional Animal Care and Use Committee approved protocol.

#### 2.1 Construction, and Purity of plasmid

Double stranded DNA fragment coding for multi characteristic opsin–II (MCO-II) was synthesized using high throughout DNA synthesizer. The restriction enzyme sites (BamH1 and Sal1) were added to the ends of the DNA fragment by PCR amplifying using the corresponding primers. Afterwards, the fragment with BamH1/Sal1 sites was inserted to the BamH1/Sal1 site of the pAAV vector. Thus, a plasmid was created comprising a CAG promoter, MCO-II and reporter (mCherry). Prior to the experiments, the plasmid was digested by restriction enzymes (BamH1 and Sal1) to verify the size and purity using 0.8% agarose gel electrophoresis.

#### 2.2 Theoretical structure prediction

Raptor-X used for predicting the 3-D structure of MCO-II. The program uses a conditional neural fields (CNF), a variant of conditional random fields and multiple template treating procedure to develop the predicted structure of MCO-II gene.

#### 2.3 Sensitizing/functionalizing gold nanorods to optical delivery beam and cells

The aspect ratio (ratio of size of short axis to long axis) of gold nanorods (GNRs) can vary from 1:1 to 1:5 to result in Surface plasmon resonance (SPR) peak varying from visible (~530 nm) to NIR (~1100 nm). Gold nanorods (diameter: 10 nm, length: 40 nm) with plasmon absorption maximum at 800 nm were used for NOD experiments. For specific membrane binding, gold nanorods were incubated with 4μM Concanavalin A at 37 °C (Overnight). The unbound ConA molecules were removed by centrifugation at 13,000 rpm for 10 min. The ConA conjugated GNR were then resuspended into 10mM HEPES buffer pH (7.2). Gold nanorods (diameter: 10 nm, length: 40 nm) with SPR maximum at 808 nm was used for NOD optimization.

#### 2.4 Patch-clamp recording setup

The MCO-II opsin is activatable over a broad visible-light spectral range (400 to 700nm). To determine the light dependent inward photocurrent, the MCO-II expressing cells were exposed to pulses (500 ms) of light with varying intensities. A single mode optical fiber coupled to a supercontinuum laser source (NKT Photonics) delivered the broadband light to the sample for optogenetic stimulation. A power meter (818-SL, Newport) was used to quantify the light intensity at the sample plane. The light pulse width was synchronized with the electrophysiology recording system, controlled by Axon Instruments Digidata system (Molecular Devices). Cells, transfected with CAG-MCO-II-mCherry were incubated with all-trans retinal (ATR, 1 μM) for 4 hours before conducting the patch clamp experiments.

The patch-clamp recording setup consists of an inverted Nikon fluorescence microscope (TS 100) platform using an amplifier system (Axon Multiclamp 700B, Molecular Devices). Micropipettes were pulled using a two-stage pipette puller (Narshinghe) to attain resistance of 3 to 5 MΩ when filled with a solution containing (in mM) 130 K-Gluoconate, 7 KCl, 2 NaCl, 1 MgCl_2_, 0.4 EGTA, 10 HEPES, 2 ATP-Mg, 0.3 GTP-Tris and 20 sucrose. The micropipette-electrode was mounted on a micromanipulator. The extracellular solution contained (in mM): 150 NaCl, 10 Glucose, 5 KCl, 2 CaCl_2_, 1 MgCl_2_ was buffered with 10 mM HEPES (pH 7.3). Photocurrents were measured while holding cells in voltage clamp at −70 mV. The electrophysiological signals from the amplifier were digitized using Digidata 1440 (Molecular devices), interfaced with patch-clamp software (Clampex, Molecular Devices). pClamp 10 software was used for data analysis.

#### 2.5 Delivery and expression of MCO-II-mCherry using NOD: HEK cells and retinal explants of rd10 mice

The HEK-293 cells were grown in static culture and maintained in Dulbecco’s modified Eagle’s medium (DMEM)/F-12 with 10% fetal bovine serum, 0.2 mg/mL streptomycin, and 200 U mL penicillin. The cultures were maintained at 37 °C in a humidified 5% CO_2_ atmosphere. Prior to the NOD experiments, the cells were incubated with ConA functionalized GNR for two hours at 37° C in a humidified 5% CO_2_ incubator. Purified plasmid (CAG-MCO-II-mCherry) was added in the cell culture medium and incubated for 15 min at 37 °C in a humidified 5% CO_2_ incubator. After 15 minutes of incubation, NOD was carried out using CW laser (800 nm).

#### 2.6 Animal preparation

B6.CXB1-Pde6b^rd10^/J mice (Jackson Laboratory, Bar Harbor, ME) were used in this study. The mice were housed in the Nanoscope technologies animal facility. All animal experimentation was conducted under local IACUC approved protocols in accordance with the ARVO statement for the Use of Animals in Ophthalmic and Vision Research. Mice were maintained on a 12:12 light cycle (lights on at 07:00). At least 6 eyes were used for in-vitro and in-vivo NOD experiments. Retinal degenerated mice (*rd10:* B6.CXB1-Pde6b^rd10^/J) were obtained from Jackson laboratory and bred in the animal facilities of the Nanoscope. Once the animals were acclimated to the animal facility for ~1 week, they were anesthetized with 90 mg/kg ketamine, 10 mg/kg xylazine acepromazine (0.5 mg/kg). In all experiments attention was paid to the ethical guidelines for investigations of experimental pain in conscious animals, and the procedures were approved by the IACUC.

#### 2.7 Retinal explant

Adult *rd10* deficient mice were sacrificed using CO_2_. The eyes of *rd10* mice were surgically removed from its muscles and remaining orbital contents and then choroid, sclera, cornea, lens, vitreous were the removed from enucleated eye. The remaining retina was incubated with fresh Neurobasal media. The RPE was gently removed from retina and retinal explants then cut into pieces using a tissue chopper. With the photoreceptor side down, the explants were placed onto sterilized petri-dishes previously coated with 5ug/dish of Poly-D-Lysine containing 1ml of Neurobasal media. The explants were incubated with ConA conjugated GNR for 2hr at 37 °C in a humidified 5% CO_2_ incubator. The CAG-MCO-II-mCherry plasmid was placed directly on the retinal surface for 15 min and explants were then exposed with 800nm CW laser for 30 sec., the GNRs are functionalized with ConA.

#### 2.8 Transduction of MCO-II-mCherry into *rd10* mouse retina

Adult (8 weeks old) *rd10* mice were treated humanely in strict compliance with IACUC on the use of animals in research. The rd10 mice (N=3) were anesthetized with a mixture of ketamine (65 mg/kg), xylazine (7.5 mg/kg), and acepromazine (0.5 mg/kg). One drop of local anesthesia (0.5% proparacaine hydrochloride) was instilled into both the eyes of the animals. Functionalized gold nanorods and MCO-II-mCherry plasmids was injected into one of the eyes by a sterilized 32-G needle of a Hamilton microsyringe inserted through the sclera into the vitreous cavity. As a negative control, the other eye was intravitreally injected with same volume of PBS. 1% Tropicamide ophthalmic solution was applied for dilating the pupil. The cornea was kept moist with a balanced salt solution during the entire surgical procedure. Schematic of experimental procedure for in-vivo NOD is shown in Fig. 4a. NOD using CW laser was carried out with the in-vivo set up (Fig. 4b). After 1 week of transfection, the animals were euthanized, and retinal tissue explanted. Using confocal microscope (Fluoview FV1000) for mCherry expression, the expression level (fluorescence intensity) of these cells were quantified. Image processing was performed by using NIH ImageJ software.

#### 2.9 Fourier domain optical coherence tomography system

FD-OCT is based on spectral domain implementation of the OCT system developed in the OCT research community^39-44^, and has been in clinical Ophthalmology practice. In the FD-OCT, reference mirror is not scanned to perform depth scan. The reference mirror is stationary. The interference signal between the reflected intensities from the reference mirror and the sample microstructures is detected with a spectrometer as a function of wavelength. The detected signal (as a function of wavelength) is then Fourier transformed to obtain intensity profile as a function of depth. FD-OCT scans the whole depth of the sample without any mechanical scanning. We performed SD-OCT imaging as described previously^45-47^. Prior to imaging, mice were anesthetized. Mice were placed on the platform of the SD-OCT and retinal thickness measurements were made.

#### 2.10 Statistics

GraphPad Prism was used to analyze data. The data were plotted as mean ± S. D. Statistical significant difference analyses were carried out by t-test. *P* < 0.05 was considered statistically significant.

## 3 Results

### 3.1. Improvement of optokinetic response in MCO-II transfected rd10

The *rd10* mice with MCO-transfected retina demonstrated increase in optokinetic response at different speed of the vertical stripes (0.1 *cpd*) at ambient light level. The basic principle of this assay is: whenever a moving pattern is presented to an animal, the (light sensing) animal will move its head as a transient corrective measure to maintain stable vision. The advantage of this method is that it does not require any previous training of the animal. Quantitative comparison of number of head movement before and after (2, 4 weeks) MCO-II transfection is shown in Suppl. Fig. 1. Quantitative comparison of number of head movement at different speed of rotation of the vertical stripes: (a) 2 rpm and (b) 8 rpm before and after MCO-II transfection. *rd10* mice with MCO-II transfection shows improved optokinetic response as reflected in the increase head movement. The average light intensity at the center of the chamber was 0.001 mW/mm^2^.

### 3.2. Nano-enhanced Optical Delivery: Principle and in-vitro optimization

The principle of Nano-enhanced Optical Delivery (NOD) is shown in Fig. 1a and 1b. The ConA conjugated gold nanorods (injected intravitreally) bind to the membrane of the targeted retinal cells. Upon illumination of a continuous wave laser of wavelength (800 nm), the enhanced photothermal properties of gold nanorods on the cell membranes resulted in localized temperature increase allowing the delivery of impermeable molecules via cell membrane. Fig. 1c shows electron microscopic image of gold nanorods used for optical enhancement of laser beam at the ends of the rods. The measured optical density of gold nano-rods with peak at 800 nm is shown in Fig.1d.

**Fig 1.**
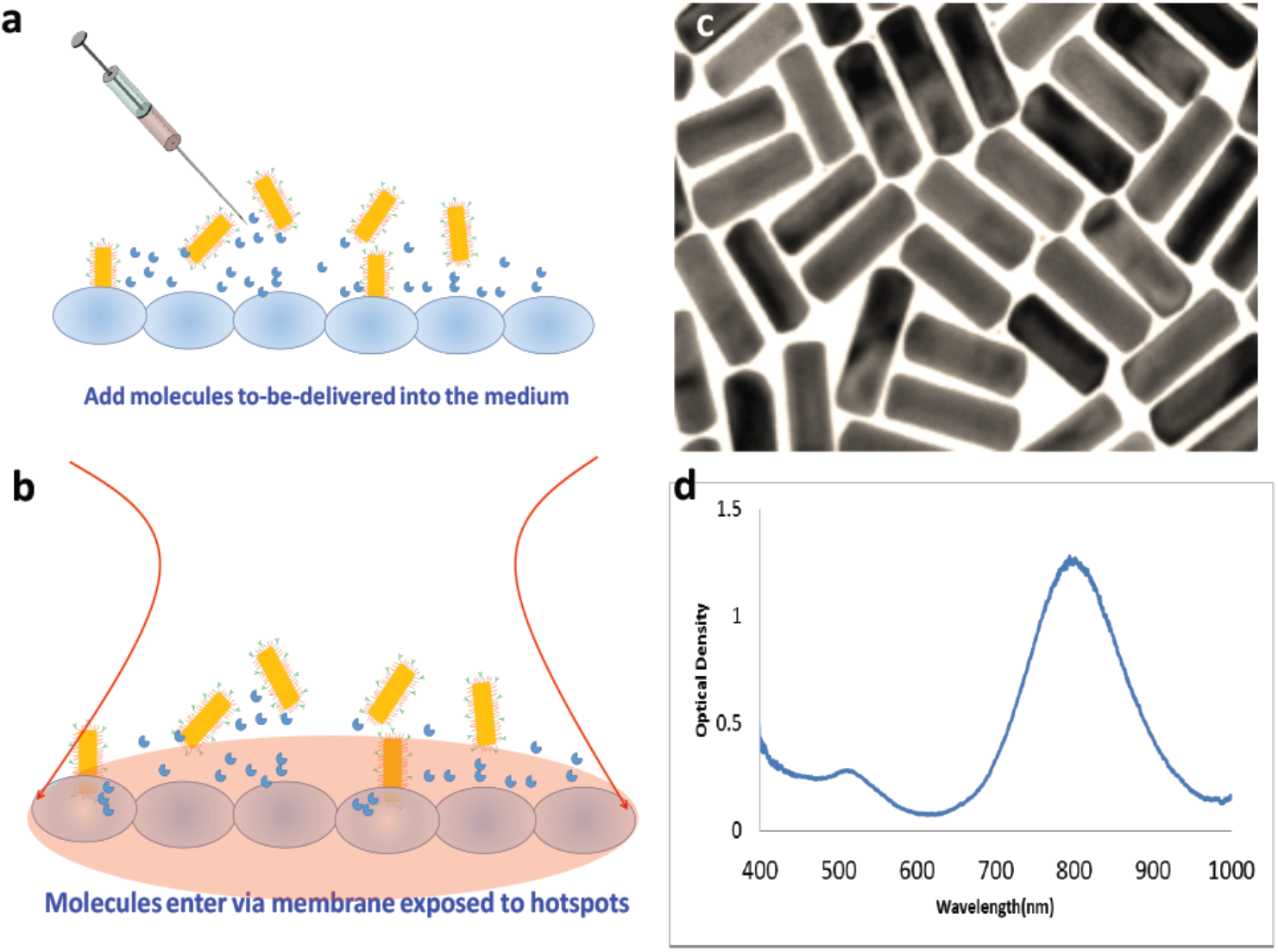
Principle of Nano-enhanced Optical Delivery (NOD). (a) The gold nanorods (injected intravitreally) bind to targeted retinal cells. (b) Upon illumination of a continuous wave laser of specific wavelength, the rods generate localized hot spots on membrane, allowing insertion of genes into the targeted (geographic atrophy) area(s) of retina. (c) Electronic microscopic image of GNRs. For binding of GNRs to cells.

NOD was used for spatially targeted delivery of ambient-light activatable multi-characteristic opsin (MCO-II) into mammalian cells. A typical circular map showing the insertion of MCO-II gene cloned in pAAV vector at the restriction sites: EcoR1 and Sal1 is depicted in Fig. 2a. The size of the vector and DNA fragment (MCO-II-mCherry) was further verified using the restriction enzymes as shown in the Fig. 2b. Fig. 2c illustrates the three-dimensional arrangement of amino acid chains of MCO-II obtained by theoretical modeling. The secondary structure prediction revealed that MCO-II opsin contains 46% of alpha helix, 17% beta sheet and 36% random coil structure. For optimization of NOD-based gene delivery, in-vitro experiments were performed for delivery of plasmids encoding for MCO-II-mCherry into HEK-293 cells. The cells were exposed for different durations to CW near-infrared laser beam with intensity of 6.4 mW/mm^2^. Fig. 2d shows a representative confocal microscopic image of HEK cells 2 days after NOD of MCO-II-mCherry plasmids in the rectangular area using 1 min exposure of CW near-infrared laser beam. The NOD-based gene delivery efficiency at different laser exposures was quantified based on the measured mCherry (reporter) fluorescence intensity of cells (averaged) in targeted areas. Fig. 2e shows the variation of mCherry fluorescence as function of NOD laser beam exposure time varying from 1 to 4 min.

**Fig 2.**
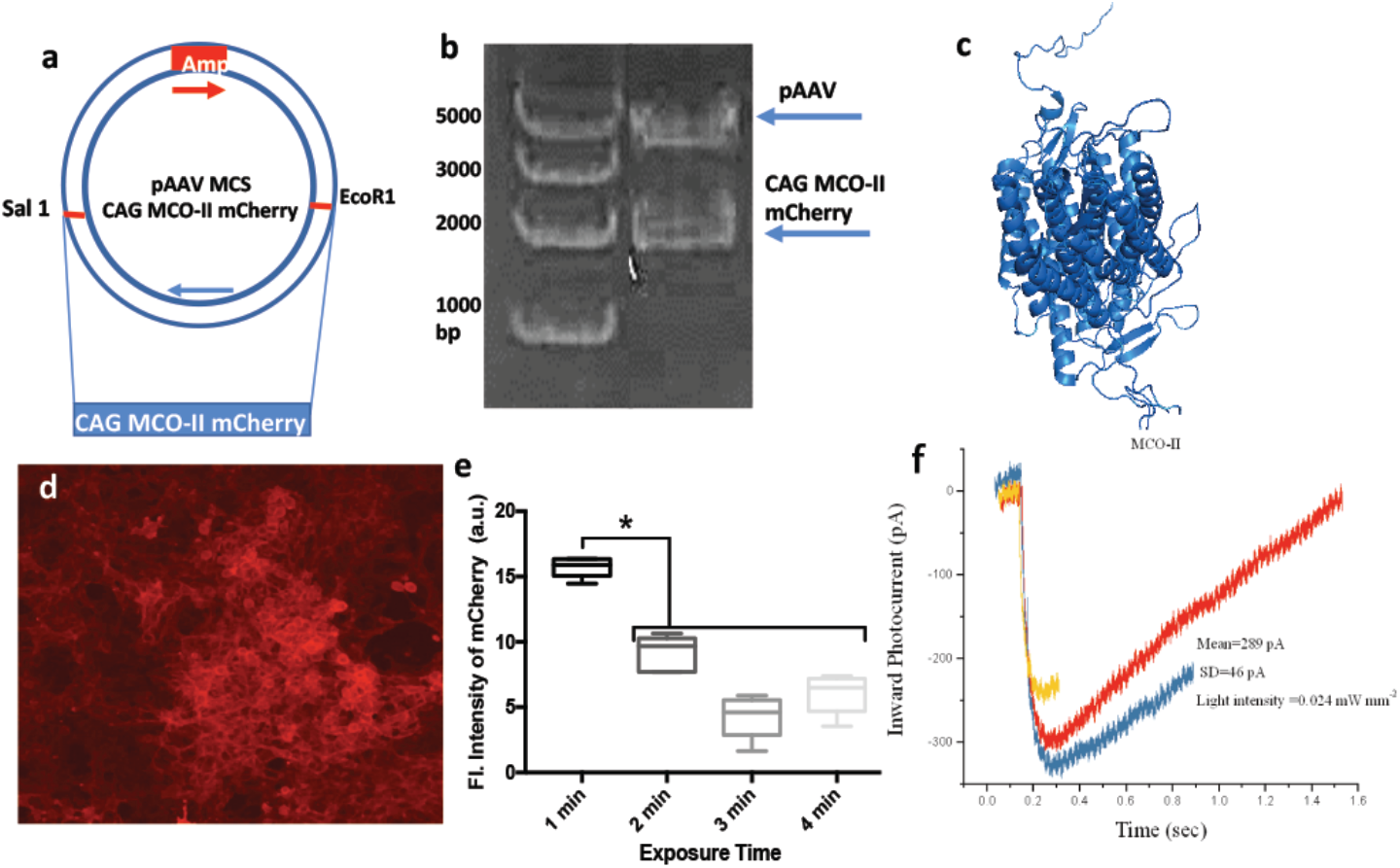
NOD optimized for delivery of Multi-Characteristic Opsin (MCO-II). (a) A typical circular map showing the insertion of MCO-II gene cloned at the restriction sites (Sal1 and EcoR1); (b) Agarose gel electrophoresis of CAG-MCO-II-mCherry gene (2915 bp) after restriction enzyme digestion with EcoR1 and Sal1; (c) Theoretical modeling of the three-dimensional arrangement of amino acid chains of MCO-II. (d) Confocal fluorescence image showing expression of MCO-II-mCherry in the HEK cells after NOD (6.4 mW/mm^2^ and 1min exposure); (e) Intrinsic fluorescence intensity of mCherry with different laser beam (800nm) exposure time, *p<0.01 between 1 min and others. (f) Inward current in MCO-II expressing cells in response to light (average intensity: 0.024 mW/mm^2^) measured by Patch-clamp electrophysiology.

To determine functioning of the MCO-II expressing cells, patch-clamp electrophysiology was carried out in the presence of white light stimulation. The representative inward current in MCO-II expressing cells in response to light (0.024 mW/mm^2^) is shown in Fig. 2f. The peak photocurrent generated in MCO-II expressing cells was found to be 300 pA. While the on-rate of induced photocurrent in MCO-II expressing cells in response to the light was in ms range, the off rate (decay of current in absence of light) was found to be significantly higher.

### 3.3. Nano-enhanced optical delivery of opsin-plasmids into retina explant

We utilized NOD parameters optimized (in HEK cells) for spatially targeted delivery of MCO-II-mCherry plasmids into retina explants. The explants were incubated for 1hr with ConA-conjugated GNRs having peak absorption at 800 nm. Confocal fluorescence image of retinal explant (Fig. 3a) shows reporter-mCherry expression in targeted area (marked by square) 2 days after exposure to CW NIR laser beam (800 nm). Fig. 3b shows cellular expression in the targeted area of the explant. The nontargeted areas (Fig. 3c) showed basal level of autofluorescence. To quantify the relative expression of mCherry in cell membrane and intracellular components, intensity profiles along line drawn across the cells are plotted (Fig. 3d). The MCO-II expression in plasma membrane was found to be significantly higher than intracellular expression. Comparison of MCO-II-mCherry expression (measured by average mCherry fluorescence intensity) in targeted versus non-targeted regions in retina explant is shown in Fig. 3e. No detectable increase in fluorescence in mCherry band was observed in non-laser exposed explants incubated with GNR and MCO-II-mCherry plasmids. Further, the explants irradiated with CW NIR laser beam and MCO-II-mCherry plasmids only (and no GNRs) did not exhibit any rise in cellular fluorescence in mCherry band.

**Fig 3.**
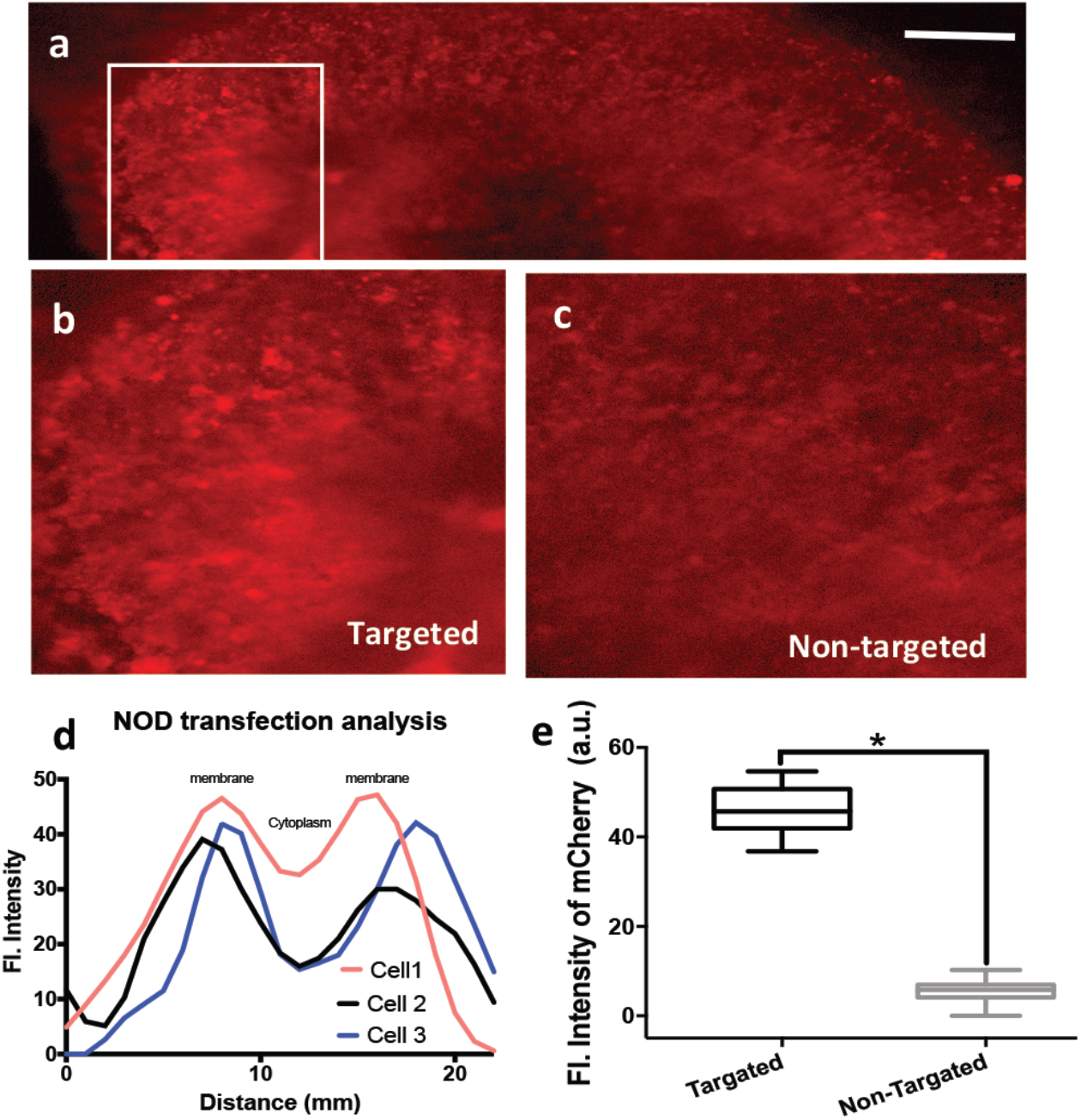
Nano-enhanced Optical delivery (NOD) of MCO-II-mCherry plasmids to retinal explants. (a) Confocal fluorescence image of retinal explant showing the expression of MCO-II in targeted area (marked by square) 2 days after exposure of CW laser (30 sec) in presence of GNR (800nm) and MCO-II plasmids, (b) Zoomed image of targeted area expressing mCherry, and (c) zoomed image of nontargeted area in an explant. (d) Intensity profile along lines drawn through the retinal cells showing preferential expression of MCO-II-mCherry after NOD in the cell membrane; (e) Comparison of MCO-II-mCherry expression in targeted versus non-targeted regions in retina explant.

### 3.4. In-vivo targeted delivery of MCO-genes to retina using NOD

After successful *in-vitro* delivery of MCO-II plasmids to retinal explants, nano-enhanced optical delivery experiments were carried out *in-vivo* using the CW NIR laser beam (800 nm) to achieve spatially targeted gene expression in the retina of *rd10* mice with photoreceptor degeneration. The experimental steps for *in-vivo* NOD into retina are shown in Fig. 4a. Fig. 4b shows the schematic of the experimental NOD setup. The eye was exposed with the CW NIR laser beam with optimized parameters (intensity: 20 mW/mm^2^, exposure time: 30 sec). 1 *μ*1 of 1: 1 mixture of ConA-functionalized GNRs and MCO-II-mCherry plasmids was injected intravitreally into *rd10* mice eye to result in final concentration of 500 ng/ml and 30 ng/ml respectively for GNRs and MCO-II-mCherry plasmids. After dilation, the CW NIR laser beam was applied for 30 seconds on the eye (Fig. 4b). Fig. 4c shows the zoomed image of an eye during the NOD laser exposure. The image of fundus with the NOD laser irradiation spot (red) is shown in Fig. 4d. 7 days after intravitreal injection and NOD, the mice were sacrificed, and retinal cup extracted and imaged using confocal fluorescence microscopy. While the non-targeted regions did not show reporter (mCherry) fluorescence (Fig. 4e), significant expression of reporter fluorescence in NOD-targeted retinal area(s) was observed (Figs. 4 f, g). Comparison of fluorescence intensity of mCherry in targeted and non-targeted regions of retina is shown in Fig. 4h. The retina of non-NOD control eye did not show any characteristic (mCherry) fluorescence. Suppl. Fig. 2 shows cellular expression of mCherry in retina extracted after in-vivo NOD of MCO-II-mCherry. Suppl. Fig. 2b shows the intensity profiles along lines drawn through retinal cells (in Supp. Fig. 2a) expressing MCO-II-mCherry after NOD. Higher intensity of mCherry fluorescence in membrane demonstrated preferred localization of MCO-II in the membrane of retinal cells as compared to cytoplasm. During intravitreal injection of MCO-II-mCherry plasmids and gold nanorods, there may be an injection related retinal damage (such as retinal detachment). Therefore, we evaluated structural integrity of retinal tissue and other ocular elements (cornea, lens) after injection (Suppl. Fig. 3 a, b). In order to characterize any damage due to NOD laser beam irradiation, the cornea and retina was imaged after NOD (Suppl. Fig. 3 c, d). Comparison of retinal thickness (No injection vs. After injection before NOD vs. After NOD) shows integrity of the retina is not compromised after NOD (Suppl. Fig. 3e).

**Fig 4.**
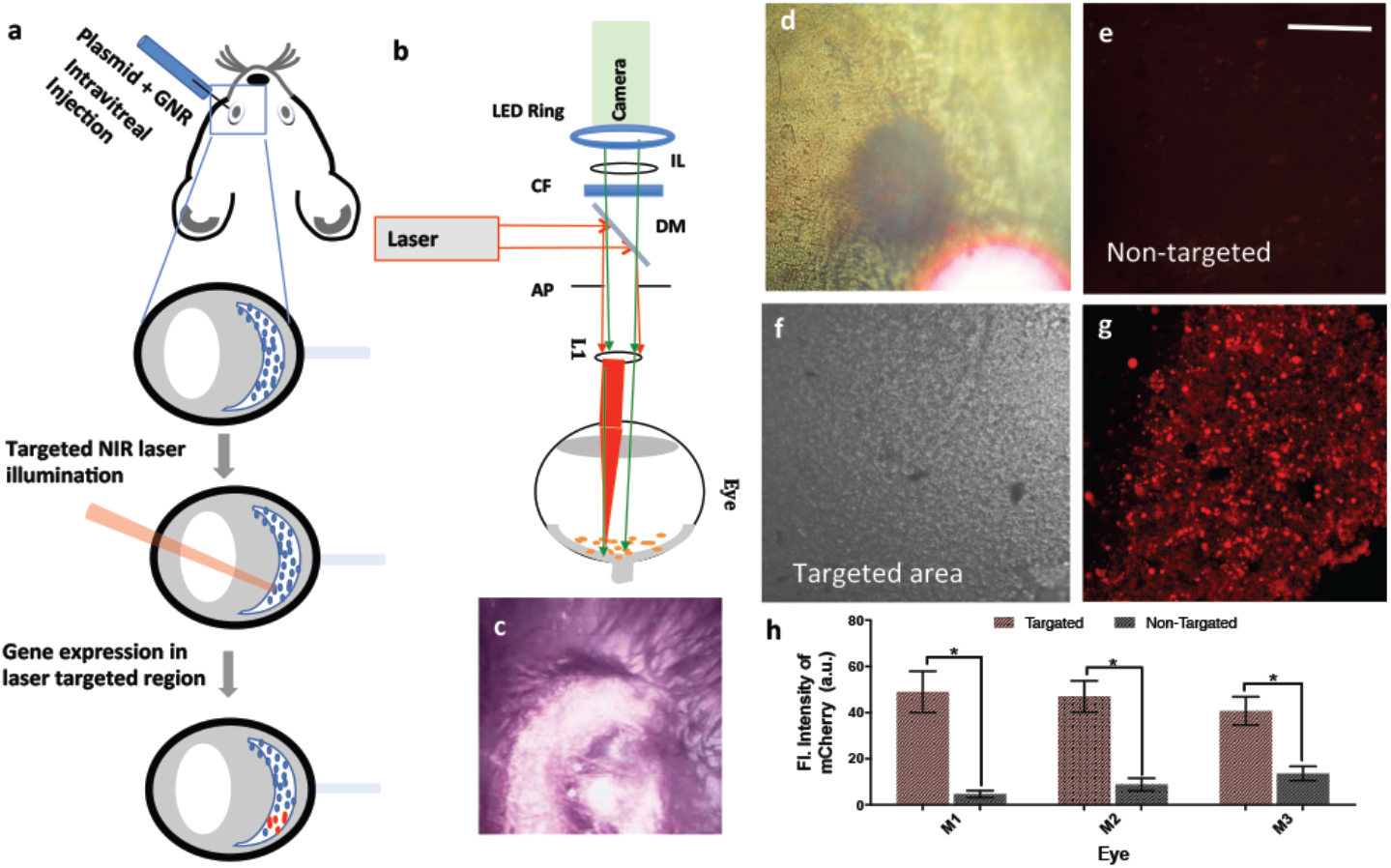
In-vivo targeted Nano-enhanced Optical delivery (NOD) of MCO-II-mCherry plasmids to retina. (a) Experimental scheme for in-vivo NOD in eye; (b) Schematic of experimental setup for in-vivo NOD. (c) Picture showing mouse eye during in-vivo NOD of the eye, injected with GNR and MCO-II plasmids; (d) In-vivo fundus image of retina during laser exposure. (e) Confocal fluorescence image of non-targeted region 7 days after NOD; (f) Bright field confocal image of the targeted region; (g) confocal fluorescence image of showing MCO-II mCherry expression 7 days after NOD; (h) Comparison of fluorescence intensity of mCherry in targeted and non-targeted regions. *p< 0.01 between targeted and non-targeted; N=10 cells/mouse.

### 3.5. MCO-II sensitized RGCs after NOD are functionally activatable by white-light

Functional response of the MCO-II expressing RGCs (in explant, delivered by NOD) toward white-light stimulation was determined by multi-electrode array electrophysiology (Fig. 5a). Different light intensities were used to stimulate MCO-II expressing RGCs. Fig. 5b shows representative spiking activities in MCO-II transfected RGCs (after NOD) with and without light. With increase in light intensity the spiking rate increased (Fig. 5c).

**Fig 5.**
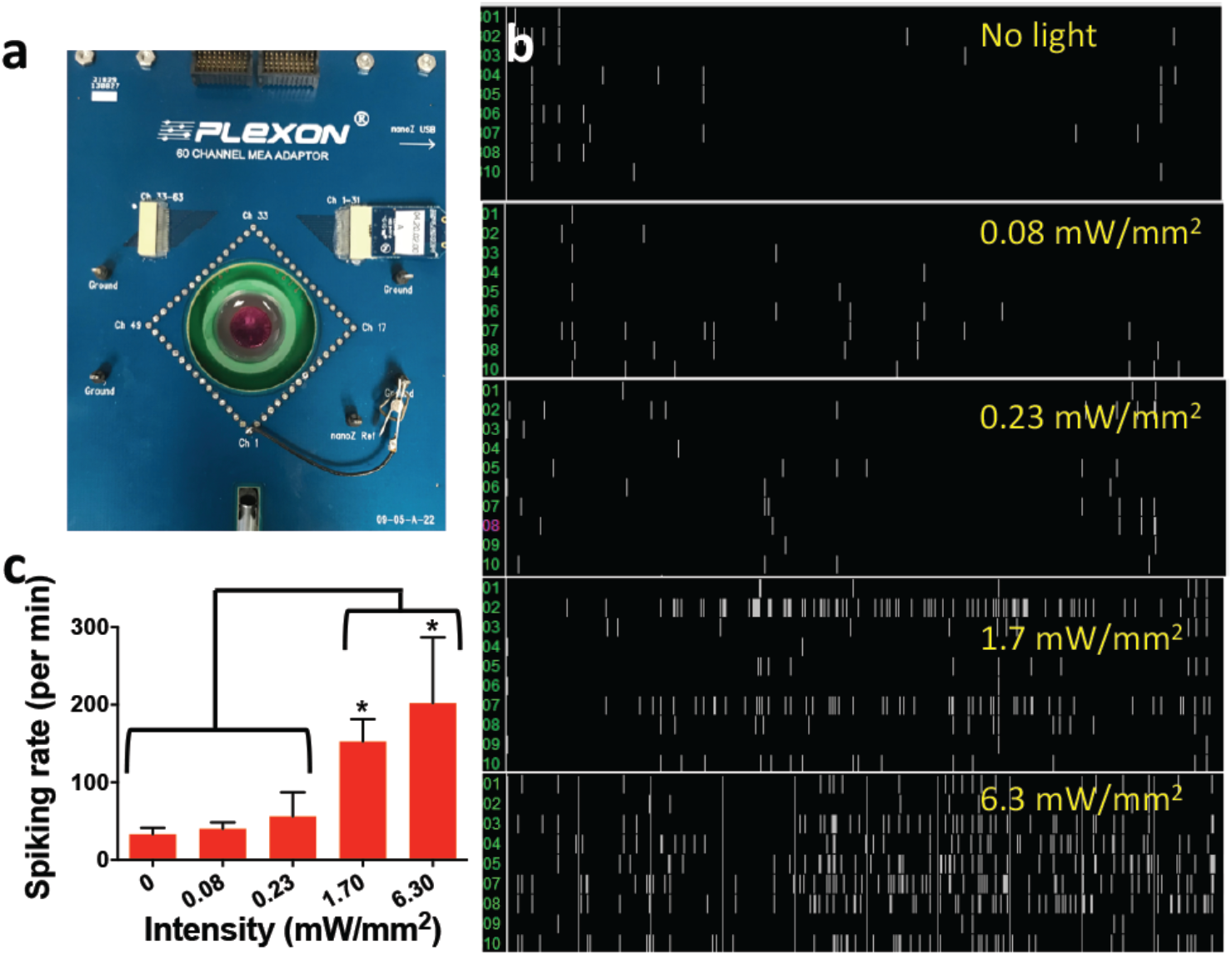
White light-activated spiking activities in MCO-II transfected RGCs in retina explant after NOD. (a) Multi electrode array chip to measure light-activated spiking in retina after MCO-II transfection using NOD. (b) White light evoked spiking activities in retina transfected with MCO-II by NOD. 1 Hz light pulses (200 ms ON, 800 ms OFF) for 60 sec. (c) Stimulation light intensity dependent spiking in retina transfected with MCO-II by NOD. *p<0.05.

## 4 Discussion

Patient-to-patient variability and time-dependent changes in spatial-distribution of retinal-degeneration demands site-specific molecular delivery for effective therapeutic outcome. For example, in case of dry-AMD, spatially targeted delivery of opsin-encoding gene is required in macula, which loses photosensitivity due to loss of photoreceptors^48-50^. With viral or other non-viral (e.g. electroporation, lipofection) method, the opsin-genes will be delivered everywhere, causing uncontrolled expression over the whole retina. This will cause complications in functioning of non-degenerated areas of retina^51^. The spatially targeted delivery into regions of degenerated retina can be achieved using NOD by shining the NIR laser beam in targeted areas. The efficient localized field enhancement by gold nanostructures^52-60^ in the safe NIR spectrum allowed *in vitro* and *in vivo* optical delivery of MCO-II-mCherry plasmid into the mammalian cells, retinal explant and targeted retinal regions of *rd10* mice. The significantly reduced white light stimulation intensity required for activating MCO-II sensitized cells will lead to ambient white light-based restoration of vision in case of retinal degenerative diseases. The NOD technology is easy to adapt in clinical setting since it puts no restriction on the maintaining the focal plane with target layer unlike focused ultrafast laser based optoporation. Further, it provides increased sensitivity of targeted cells with no detectable collateral damage. In addition, nano-enhancement near gold nanorods (used in NOD) allows use of low power, compact, less expensive and easy-to-use lasers.

Further, in case of loss of opsin expression in retina or development of new atrophies, it will be necessary to re-induce the opsin-encoding genes. The viral vectors have limitations in subsequent gene delivery in humans due to immunorejection and are unable to substantially re-induce transgene expression by reinjection in rodents^61^. In contrast, NOD has the ability for reinduction of genes to retina, providing an important advantage over viral vectors. Furthermore, the current state-of-the-art method for preclinical and clinical ocular gene therapy uses viral vectors, especially AAV vectors, which have selective tropism for certain retinal cell types. There often is a need to target multiple retinal cell types. For example, in glaucoma and ischemic retinopathy one need to target RGCs, microglia, astrocytes, and invading macrophages^62-63^. By varying shape of functionalized field-enhancing gold nanorods that binds to targeted cell types, tuning the NIR laser wavelength will allow wavelength-selective enhancement near specific cells and thus, selective delivery.

In NOD method, the contrast in temperature rise in laser-irradiated nanorod-attached cells at nano-hotspots is significant enough to allow site-specific delivery of large (MCO-II-mCherry) plasmids. The NOD approach based on CW NIR laser beam is found to be minimally invasive with no detectable collateral damage to retina or other ocular elements as measured by OCT. Further, the gold nanorods have very low cytotoxicity and have been used in clinical trials. In case of rapid movement of eye in clinical practice, it may be advantageous to use a spatially sculpted NIR beam to match the shape of the region(s) of interest in the tissue requiring targeted molecular delivery by NOD. Shaping the NIR laser beam for NOD can be achieved by use of digital micromirror device (DMD) or spatial light modulator (SLM) to fit the targeted regions (e.g., geographic atrophies of retina in degenerative diseases) so that the therapeutic molecules (e.g. genes) can be delivered in a high throughput manner.

## 5 Conclusions

The *in-vitro* and *in-vivo* results, presented here using NOD, clearly demonstrate *in-vivo* gene delivery and functional cellular expression in targeted regions of retina using CW NIR laser beam without compromising structural integrity of the eye. Our results show that significant photocurrent can be reliably generated in MCO-II sensitized cells at white light intensity level close to ambient light condition. We believe our method of non-viral delivery of gene encoding opsins will pave the way for clinical applications in treatment of patients with retinal dystrophies.

## Author Contributions

The manuscript was written through contributions of all authors. / All authors have given approval to the final version of the manuscript.

## Acknowledgement

The authors would like to thank Dr. Sanjay Pradhan (Nanoscope) for animal care and Harvey Wiggins (Plexon Inc) for MEA equipment.

## Supplemental Information Nano-enhanced optical gene delivery to retinal degenerated mice

**Suppl. Fig. 1.**
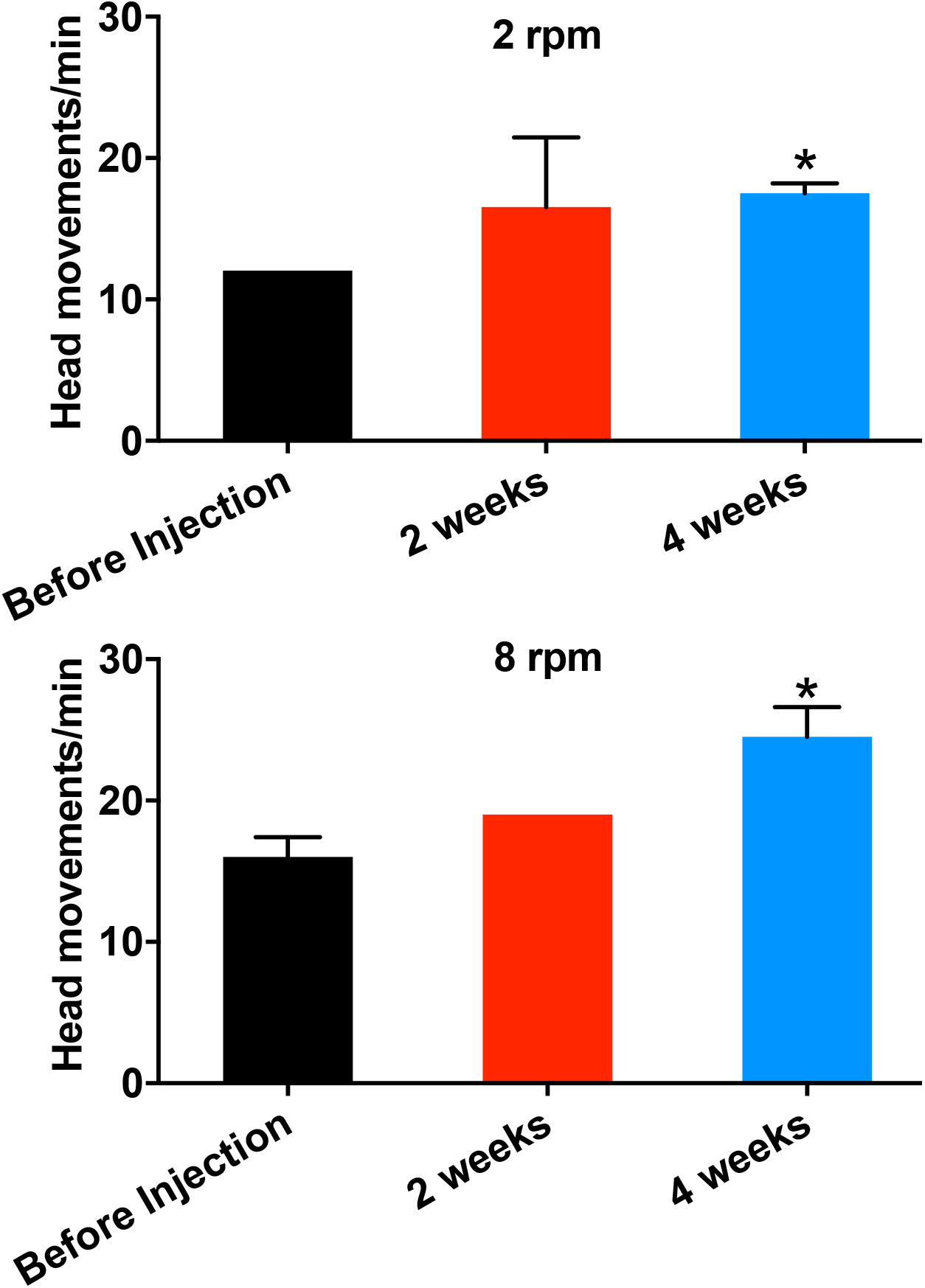
Improvement of optokinetic response in MCO-II transfected rd10 mice at ambient light level. Quantitative comparison of number of head movement at different speed of rotation of the vertical stripes: (Top) 2 rpm and (Bottom) 8 rpm before and after MCO-II transfection. *rd10* mice with MCO-II transfection shows improved optokinetic response as reflected in the increase head movement. The average light intensity at the center of the chamber was 0.001 mW/mm^2^.

**Suppl. Fig. 2.**
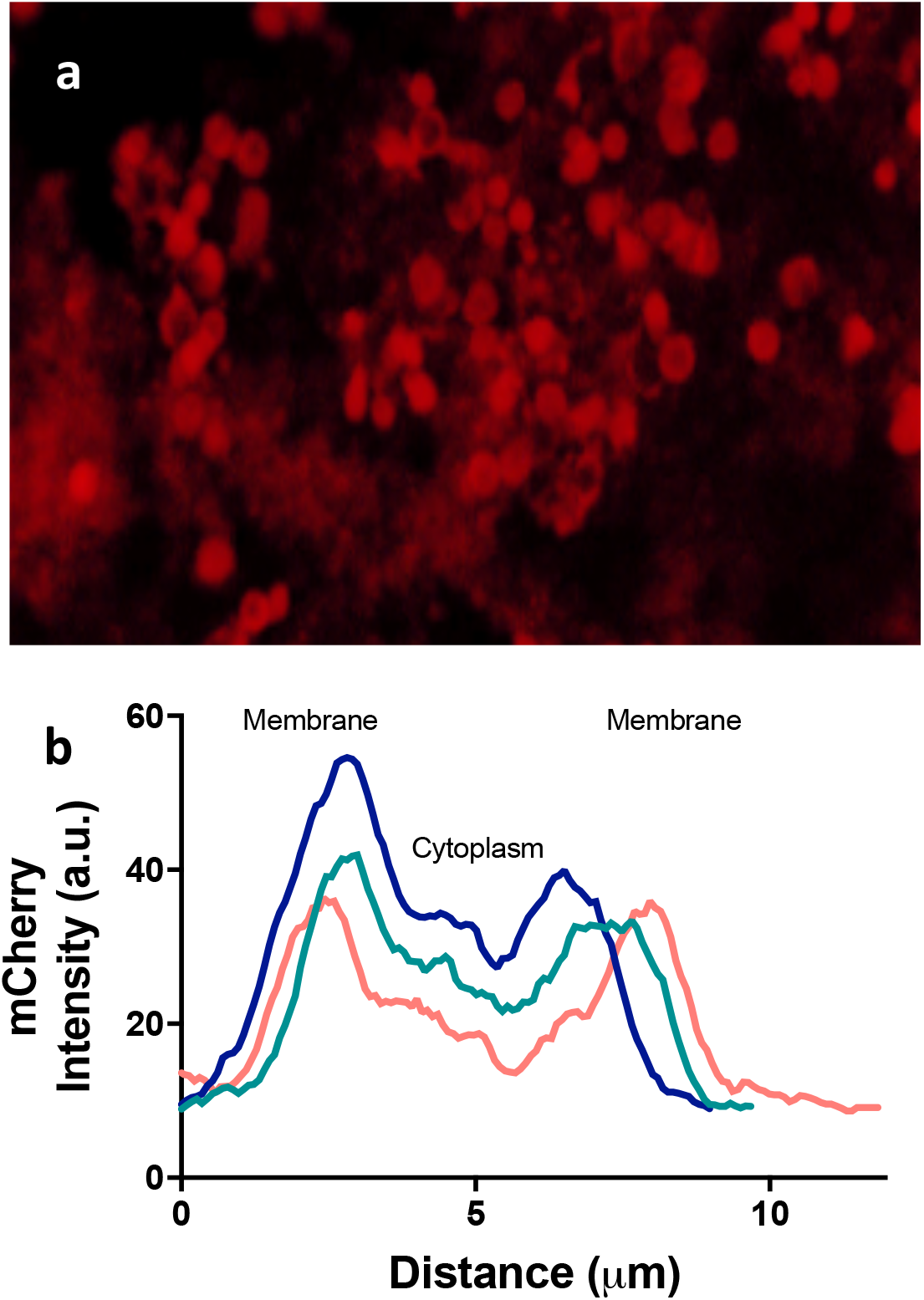
(a) Confocal image showing expression of MCO-II in the retinal cells after ~1 wk of NOD. (b) Intensity profile along lines drawn through the cells expressing MCO-II-mCherry after NOD.

**Suppl. Fig. 3.**
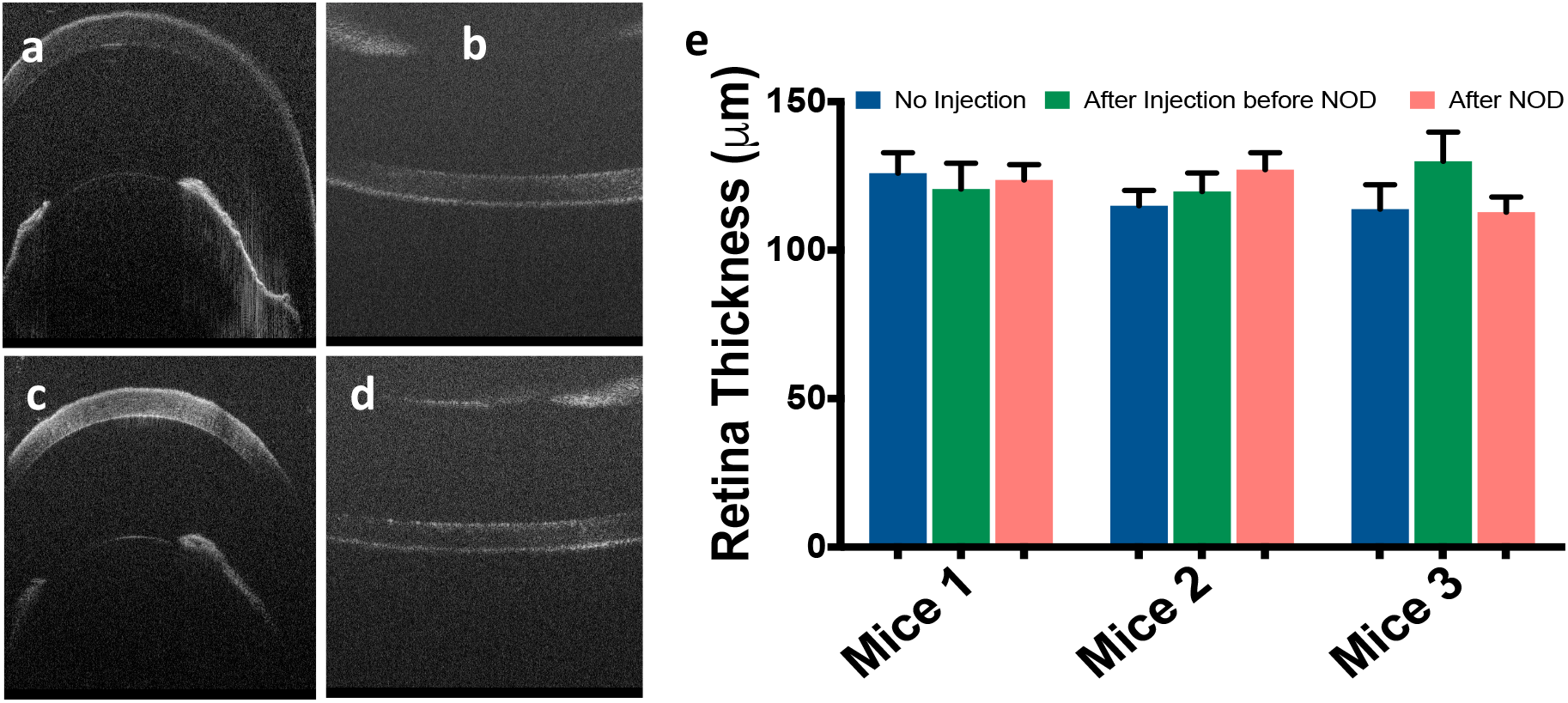
(a) Monitoring changes in cornea and retina using SDOCT after NOD. OCT image of (a) cornea and lens (and iris) and (b) retina before optical delivery. OCT image of (c) cornea and lens (and iris) and (d) retina of same animal after optical gene delivery to retina. (e) Comparison of retinal thickness (No injection vs. After injection before NOD vs After NOD).

